# Long-term cardiovascular re-programming by short-term perinatal exposure to nicotine‘s main metabolite cotinine

**DOI:** 10.1101/193003

**Authors:** Stefano Bastianini, Viviana Lo Martire, Alessandro Silvani, Giovanna Zoccoli, Chiara Berteotti, Hugo Lagercrantz, Anders Arner, Gary Cohen

**Author notes:** **Corresponding Author:** Gary Cohen Centre for Sleep Health and Research & Sleep Investigation Laboratory Royal North Shore Hospital, Reserve Road, St Leonards, NSW 2067, Sydney, Australia **Email:** **Email:**.

## Abstract

**Background:** Cotinine - a nicotine by-product and biomarker of passive perinatal tobacco smoke exposure - is historically considered to lack significant health effects. We challenged this notion and sought “proof-of-concept” evidence of the adverse developmental potential of exposure to this substance at real-life levels.

**Methods:** Pregnant C57 mice drank nicotine or cotinine-laced water for 6wks from conception (N_PRE_ = 2% saccharin+100μg nicotine/ml; C_PRE_ = 2% saccharin + 10μg cotinine/ml) or for 3wks after birth (C_POST_ = 2% saccharin + 30μg cotinine/ml). Controls drank 2% saccharin (CTRL). At 17±1weeks male pups (CTRL n=6; C_POST_ n=6; C_PRE_ n=8; N_PRE_ n=9) were instrumented for EEG and blood pressure (BP) telemetry. We evaluated (i) cardiovascular control during sleep (at rest / during stress); (ii) arterial reactivity ex vivo; (iii) expression of genes involved in arterial constriction/dilation.

**Results:** Blood cotinine levels (ELISA) recapitulated passive smoker mothers-infants. Pups exposed only to cotinine exhibited (i) mild bradycardia - hypotension at rest (p<0.001); (ii) attenuated (C_PRE_, p<0.0001) or reverse (C_POST_; p<0.0001) BP reactivity to asphyxia; (iii) pronounced adrenergic hypo-contractility (p<0.0003), low Protein Kinase C (p<0.001) and elevated adrenergic receptor mRNA (p<0.05) (all drug-treated arteries). N_PRE_ pups also exhibited endothelium-mediated dysfunction.

**Conclusions:** Cotinine has subtle, enduring developmental consequences. Some cardiovascular effects of nicotine can plausibly arise via conversion to cotinine. Low-level exposure to this metabolite may pose unrecognized perinatal risks. Adults must avoid inadvertently exposing a fetus or infant to cotinine as well as nicotine.

## Introduction

A pregnant passive smoker ingests cumulatively far less of the semi-volatile toxin nicotine than a pregnant active smoker, yet offspring of the former exhibit a disproportionate anomaly in postural blood pressure compensation, a finding difficult to reconcile with exposure to nicotine *per se* notwithstanding its potency as a cardiovascular toxin (1). Most (80%) inhaled or ingested nicotine is rapidly broken down in the liver to cotinine. Cotinine -a minor structural variant (and anagram) of nicotine – is historically considered relatively “safe-inert” i.e. lacking significant pharmacological, cardiovascular or other side effects even at blood levels 10-fold higher than attained by active smoking (2, 3). Cotinine eventually passes out of the body in urine, but compared to its parent compound blood levels rise far higher (>5-10 fold) (4, 5) and stay high for much longer by virtue of slow body elimination (half life = 19-24hr vs. 2hr for nicotine) (6). Cotinine penetrates tissues more extensively than nicotine, and also lowers the heart rate (HR) and blood pressure (BP) of adults - effects opposite to nicotine itself (7-9). It has a different mode of action to nicotine: evidence suggests cotinine is a positive allosteric modulator of the naturally occurring acetylcholine receptor (nAChR) hence potentiates the responsiveness of nAChRs to, and sensitizes tissue to the effects of, natural ligands (acetylcholine, choline) and agonists like nicotine (10, 11). On this basis, it might reasonably be expected that prolonged perinatal exposure will have some consequences (12). Here, we undertook a systematic study – the first of its kind – to differentiate long-term effects of early-life exposure to cotinine from those of its parent alkaloid nicotine.

To explore the effects of low-level, human maternal-fetal (*in utero*, via the placenta) and mother-to-newborn (*ex utero*, via breast milk) exposure to cotinine or nicotine we developed a novel mouse model. Our aim was to simulate, as realistically as possible, cotinine’s potential perinatal actions. In-principle proof that cotinine is more than just an inert biomarker of recent exposure to nicotine and/or second hand smoke (SHS) has broad public health implications, in terms of global efforts to eradicate *all* avoidable mother-infant SHS exposure (1), as well as perhaps for pregnant smokers who contemplate using nicotine replacement therapy (NRT) to quit. Based on World Health Organization estimates of the prevalence of SHS exposure, up to 40% of infants and children globally receive a stubbornly, alarmingly high ongoing cotinine load via indirect and/or direct ingestion of nicotine exhaled by smokers (13).

Recommending NRT to pregnant or breast-feeding women as “safe and effective”, and calls to raise the current dosage regimen because it “does not work well enough” (14, 15) may also need to be reconsidered if NRT’s main by-product cotinine can be shown to have *unrecognized* adverse developmental side effects.

## Methods

The experiments were approved by the Karolinska and Bologna University ethics committees for animal experimentation, and were conducted according to the principles expressed in the Declaration of Helsinki.

## Mice and Drugs

C57Bl/6J dams (3-4/cage) drank water laced with 2% Saccharin (CTRL), or 2% Saccharin+100μg/ml nicotine freebase (N_PRE_; Sigma Aldrich), or 2% Saccharin+10μg/ml cotinine (C_PRE_; Sigma Aldrich, C5923) for 3 weeks prior to and 6 weeks after mating (i.e. exposure was continuous, commencing at conception and ceasing at or shortly after weaning); cotinine at this dose matched the blood level of N_PRE_ dams (Figure 1A). An additional cohort (C_POST_) drank 2% Saccharin+30μg/ml cotinine for 3 weeks exclusively after birth until weaning, to achieve *de novo* postnatal-only exposure equivalent to that of the other litters. Drug-laced drinking water was refreshed twice weekly.

**Figure 1.**
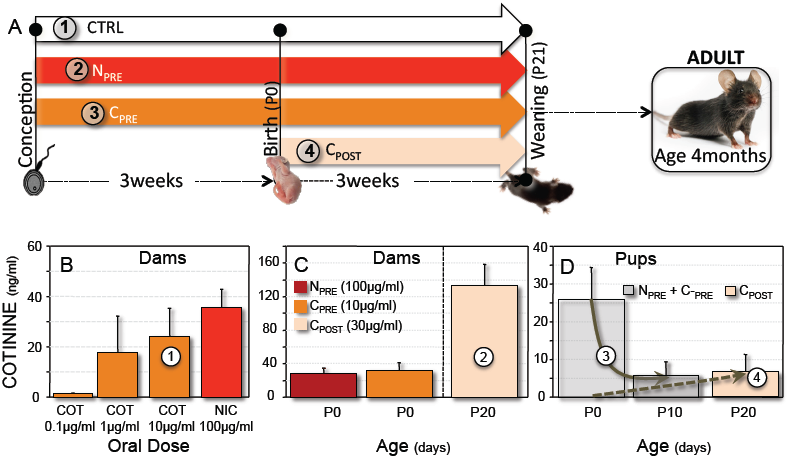
Plasma Cotinine (ELISA). The experimental protocol is summarized in the upper panel (**A**); note pups were tested 4months after cessation of all drug exposures regimens. Inital trials established the oral cotinine dose needed to roughly match the plasma cotinine level of dams given low-dose nicotine (**1 in panel B**). Dams given a higher oral cotinine dose for 3wks *after* giving birth had >3-fold higher blood levels vs. other dams (**2 in panel C**). Cotinine in blood from a subset of newborns exposed from conception plummeted after birth (**3 in panel D**, arrow; p=0.015) presumably due to loss of placental transfer. Litters exposed exclusively to cotinine sequestered in milk (C_POST_) had blood levels at weaning equivalent to the postnatal minimum of both litters exposed from conception (**4 in panel D**). Means ± SEM.

## Experimental protocol

Tail blood (100μl) was collected from anesthetized dams 3wks after drug exposure started and again immediately post-partum. Some N_PRE_ and C_PRE_ pups were sacrificed on the day of birth, or on postnatal day 10 for pooled trunk / heart blood collection; tail blood (100μl) from anesthetized C_POST_ pups was collected on postnatal day 20 i.e. just prior to weaning. Blood was centrifuged (2000rpm x 10min at 4°C) and plasma kept at -80°C until assayed (Mouse/Rat Cotinine ELISA, Calbiotech). Weaned pups were kept under a 12:12-h light–dark cycle (lights on at 9AM) at 25°C with free access to drug-free drinking water and food (4RF21; Mucedola, Settimo Milanese, Italy).

## Phenotyping

At 17±1weeks male pups (to avoid confounding effects of oestrus) from CTRL (n=6), C_POST_ (n=6), C_PRE_ (n=8), and N_PRE_ (n=9) dams were anesthetized (1.8-2.4% isoflurane-O_2_ + Carprofen 0.1mg sc; Pfizer Italy) and implanted with electroencephalography (EEG) / neck electromyography (EMG) electrodes, and a telemetric arterial blood pressure (BP) transducer (TA11-PAC10, DSI, Tilburg, Netherlands) (16). The transducer tip was advanced via the femoral artery into the abdominal aorta just caudal to the the renal arteries. After ∼2wks recovery each mouse pup underwent a standard EEG/EMG/BP morning recording in a whole-body plethysmograph consisting of 5hr habituation (chamber continuously purged with air) followed immediately by ∼2hr intermittent mild asphyxia during sleep (chamber purged with 5cycles of 15min 13%O_2_+2%CO_2_–10min air). Asphyxia simulates (over an extended time course) mild sleep apnea and helps to unmask underlying, “silent” dysfunction (17). Gas exiting the plethysmograph was continuously sampled (ML205 Gas analyser, ADInstruments).

## Vascular Reactivity

Once phenotyping was complete, aortas from the arch to just rostral to the renal arteries were harvested under deep anesthesia, stored at 4°C in CO_2_-bubbled Krebs-HEPES (18) and despatched overnight from Bologna to Stockholm (1 of 6 batches of vessels inadvertently lost in transit). The following morning 2-3 rings ∼1mm long cut sequentially in a caudo-rostral direction were mounted on tungsten pins (620M Multi Wire Myograph, Aarhus, Denmark) suspended in Krebs buffer bubbled with 5%CO_2_-95%O_2_ at 37°C (refreshed each 15min). Rings were gently stretched co-axially (∼5mN, optimal resting tension for 22–28g mice), stabilized for 1hr, then subjected to: (i) 2x80mM KCl each 30min to measure contractile force (mN) and tension corrected for segment length (mN/mm); (ii) cumulative doses of the adrenergic agonist phenylephrine (PHE) to assess reactivity; (iii) pre-contraction with 10^-6^M PHE then stepwise doses of acetylcholine (ACh) to test endothelium-dependent relaxation; (iv) 10^-4^M sodium nitroprusside (SNP) to elicit full, endothelium-independent relaxation.

## Quantitative Real-Time Polymerase Chain Reaction (qPCR) mRNA expression

Carotid arteries harvested in tandem with aortas were kept at -80°C until RNA extraction (RNeasy Plus Universal Mini Kit, Qiagen, Italy). From 60ng/sample converted to cDNA we compared the expression of 7 genes involved in smooth muscle contraction-relaxation (GoTaq® 2-Step RT-qPCR System, Promega, Sweden), using forward and reverse primers designed to span the exons of each gene via web-based software (http://www.kegg.jp/kegg-bin/highlight_pathway?scale=1.0&map=map04270&keyword=smooth%20muscle%20contraction; Table 2). The mRNA of the 7 targets - the adrenergic α1D receptor (ADR), L-type high voltage-gated Ca^2+^ channel α1 subtype (HVCC), Ras homologA GTPase (RhoA), protein kinase C (PKC), myosin light chain kinase (MLCK), Rho kinase (ROCK), myosin phosphatase inhibitor protein (CPI-17) - was expressed relative to a reference gene (β-actin) known not to be intrinsically affected by nicotine at the dose used (19). We used an i-Cycler IQ5 (Bio-rad) and annealing T=60°C, except for HVCC, CPI-17 and PKC (T=57°C to reduce non-specific amplication); specific amplification for each primer pair was assessed from the melting curves at the end of each assay.

## Analysis and Statistics

Sleep state was scored as described previously (16). Movement (MVT) was artifact ≥1s (17). We calculated beat-to-beat systolic (SBP), diastolic (DBP), pulse pressure (PP) and heart rate (HR), instantaneous breath rate (f) and weight-adjusted tidal (V_T_) and minute (V_E_) volumes (Labchart v7, AD Instruments)(16). The initial two asphyxia cycles invariably disrupted sleep so were excluded. The non-rapid eye movement (”quiet”) sleep steady-state response (SSR) was the 11^th^-15^th^ min of the 3^rd^–5^th^ asphyxia cycles inclusive (excluding MVTs and epochs of other states). The BP of one pup was excluded due to telemetry malfunction. We used a 2-factor ANOVA with drug exposure and gas (air/asphyxia) as factors, then Bonferroni-adjusted t-tests if the omnibus p<0.05. Cotinine was analysed non-parametrically (Mann-Whitney / Kruskall Wallis test). Since cotinine’s toxicity is a fraction of nicotine’s (2, 10) and C_POST_ mice received the lowest *cumulative* cotinine exposure, cumulative dosage was qualitatively expressed as CTRL<C_POST_<C_PRE_<N_PRE_ (x-axis Figure 4B). From each vascular dose-response curve fitted by nonlinear regression (GraphPad Prism V6, San Diego, CA) we determined-log concentration at half-maximal response (EC50). ACh-induced relaxation is shown % pre-contraction. The mRNA ratios are shown relative to β-actin (ΔΔCt method) (20). Data are means ± SEM.

## Results

Litter size and birth weight were comparable across groups, although at age ∼4months C_PRE_ males were marginally heavier than any other litter (Table 1).

## Cotinine transfer to pups – high before, low after birth

The mean cotinine level of N_PRE_ dams was less than reported for the same oral regimen (21, 22) but matched that of C_PRE_ dams (Figure 1B). Cotinine in fetuses exposed *in utero* fell by ∼80% after birth presumably due to elimination of placental transfer (low-level exposure continued via milk; Figure 1D). As anticipated, plasma cotinine levels were highest in C_POST_ dams, but their pups received (via milk) *de novo* exposure roughly equivalent to the postnatal minimum of both other groups (Figure 1D). Cotinine was undectable in CTRL dams or pups.

## Sleep – wake behaviour: unchanged by perinatal drug exposure

Quiet sleep consolidated with each bout of asphyxia (Figure 2A-B) and prevailed during cycles 3-5 (Figure 2C). Total and average duration of MVTs disrupting sleep cycles 3-5 (Figure 2 D-E) was comparable across groups, indicating a key protective-defensive response to an impending threat was essentially intact after exposure to nicotine or its metabolite.

**Figure 2.**
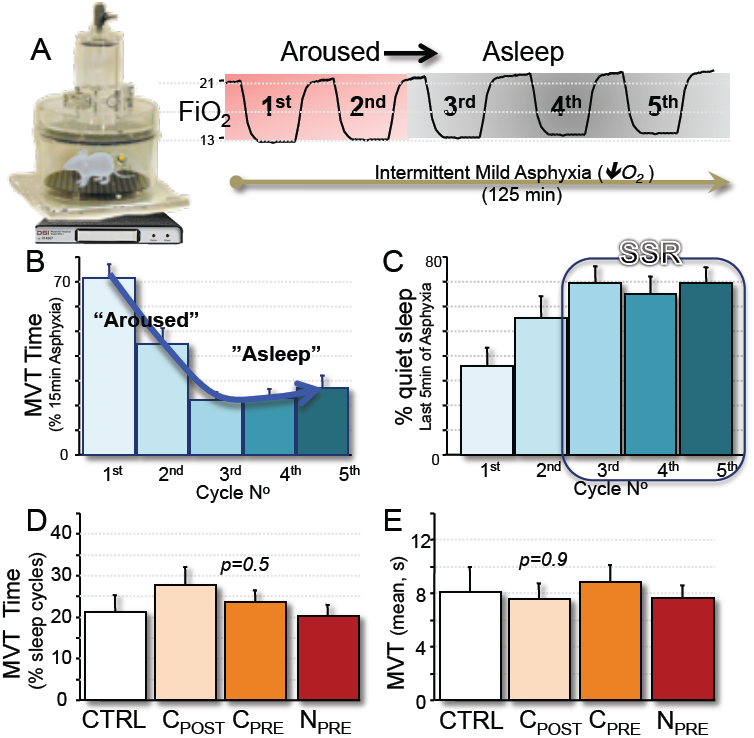
Cotinine – no effect on sleep and arousal. The protocol is summarized (**A**). Body movements lasting ≥1s (MVT) initially disrupted quiet sleep but were shorter and less frequent with each asphyxia cycle (**B**; blue arrow). Non-rapid eye movement (“quiet”) sleep predominated after the 2^nd^ bout of asphyxia (**C**; % final 5min of asphyxia cycles 3-5, groups combined for clarity), from which the steady-state response (SSR) to asphyxia was determined. Total (**D**; % cycles 3-5) and average MVT duration (**E**) indicate a key protective reflex that triggers / initiates arousal from sleep was functionally intact in all drug-exposed litters. Means±SEM

## Cotinine: devoid of respiratory side effects but slows the heart and lowers BP

Exposure to nicotine (but not cotinine) produced a persistent, underlying hyperpnea (tidal and minute volumes consistently ∼17% and ∼6% greater, respectively, vs. all other groups; Table 1). Asphyxia elicited the same relative increase from all pups (Figure 3A-B; SSR ∼7%/group; p=0.9) due to “rapid-shallow” breathing (Figure 3D). Cotinine-exposed mice were mildy but significantly bradycardic-hypotensive at rest, but nicotine lowered only pulse pressure (narrower systolic-diastolic gap; Table 1). The typical circulatory response to asphyxia – a reciprocal rise in HR / fall in BP (Figure 4A) – was evident in nicotine-treated but not cotinine-treated mice: the latter’s BP response was weak (C_PRE_) or absent (C_POST_) (Figure 4B1-B2). In short, since cotinine’s effects on blood pressure control were opposite to nicotine’s the dose-effect curve was non-linear / an inverted U-shape ((1)Figure 4B1-B2).

**Figure 3.**
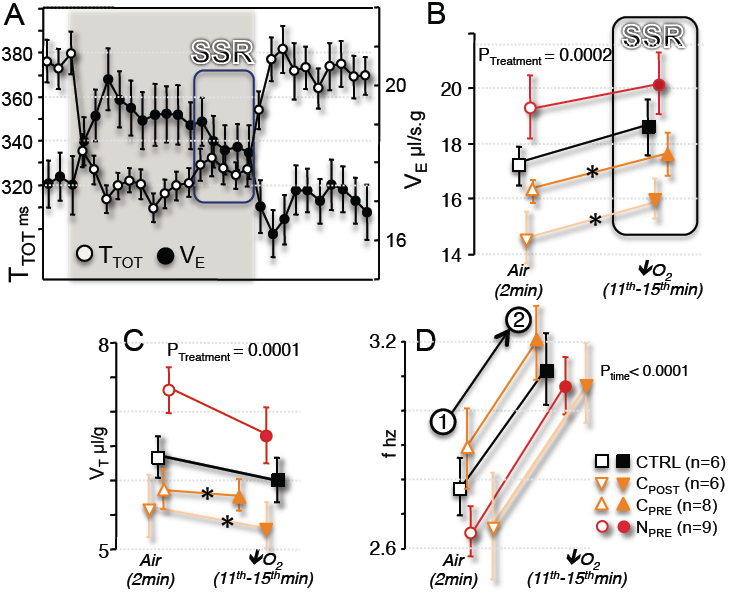
Cotinine is devoid of respiratory side effects. Breath duration (T_TOT_; **A**) and weight-corrected minute volume (**A-B**; V_E_=V_T_ x f) and tidal volume (V**_T_**; **C**), breath rate (f; **D**), and the net steady-state response to asphyxia (i.e. the 11-15^th^min of the 3^rd^-5^th^ quiet cycles = SSR) are shown. Conclusions: (i) the rate of increase in V_E_ during asphyxia was equivalent for all groups (i.e. slope of the SSR; **B**; p=0.9) due to “rapid-shallow” breathing by all (arrow joining 1-2 in **D**), however (ii) the average V_T_ and V_E_ of N_PRE_ pups was consistently greater vs. C_POST_ and C_PRE_ groups (upward displacement of lines in **B**-**C**; *p<0.05 vs. N_PRE_).

**Figure 4.**
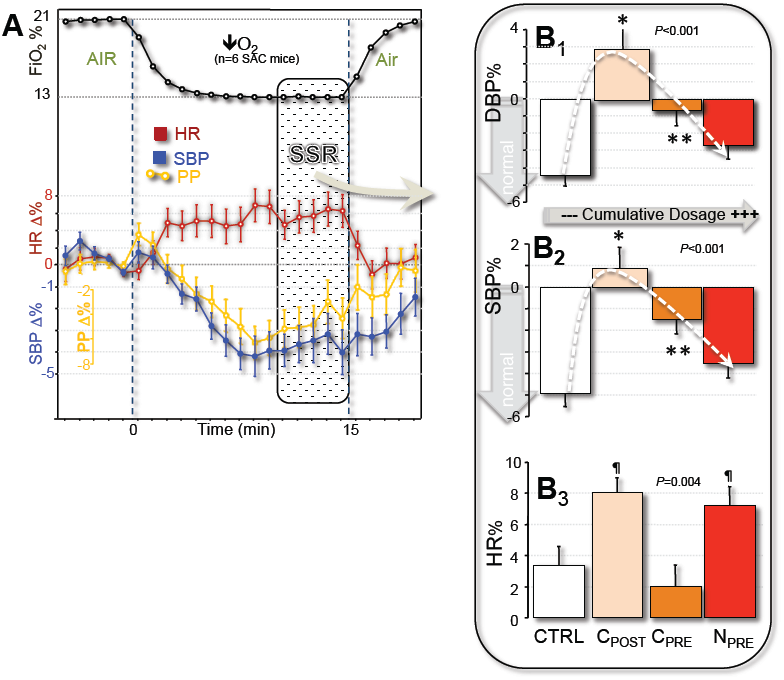
Cotinine’s effect on blood pressure is opposite to nicotine’s. The systolic (SBP), diastolic (DBP), pulse pressure (PP), and heart rate (HR) response to repetitive mild asphyxia (% change from preceding 2min air in **A**; quiet sleep asphyxia cycles 3-5 combined) and the peak quiet sleep steady-state response to asphyxia (SSR=11^th^– 15^th^ min mean; **B1-B3**). Comparative cumulative drug dosage is shown qualitatively on the x-axis (*Methods*). **Conclusions**: (i) HR normally rose and BP fell slightly during asphyxia; (ii) postnatal cotinine exposure (C_POST_) weakened BP control more than either of the other drug exposure regimens, the inflexion point of the dose-effect curves (**B1-B2**) most likely signifying some adaptive response to prolonged gestational drug exposure. Means ± SEM. *p<0.05 vs. CTRL; **p<0.05 vs. CTRL and vs. C_POST_.

## Cotinine weakens adrenergic vasoconstriction, nicotine impairs vasodilation

KCl-induced depolarization-Ca^2+^ influx-smooth muscle contraction revealed that vessels from N_PRE_ mice generated intrinsically greater tension (Figure 5A). Activation by PHE showed comparable adrenergic hypo-contractility of N_PRE_ and C_POST_ vessels (right-shift in dose-response curves, ∼two-fold higher EC50; p<0.0003; Figure 5B; similar but non-significant shift for C_PRE_). Pre-contracted CTRL and cotinine-treated but not N_PRE_ vessels relaxed normally with ACh (Figure 5C), whilst SNP completely relaxed all vessels. In short: endothelial dysfunction was attributed to nicotine but vascular hypo-contractility to cotinine.

**Figure 5.**
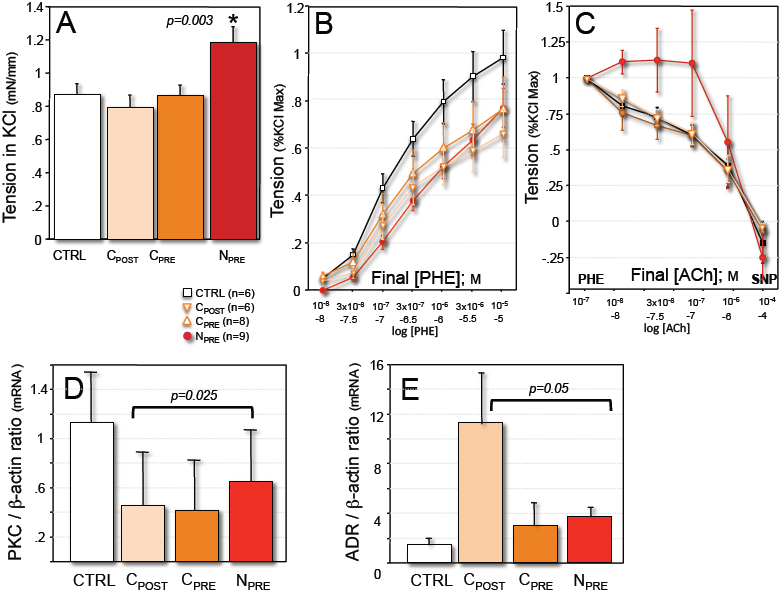
Vascular reactivity and gene expression. Nicotine-treated vessels contracted with more force in response to direct (K^+^-mediated) depolarization-Ca^2+^ influx (**A**). Reactivity to phenylephrine (PHE) revealed pronounced adrenergic hypo-contractility of nicotine *and* cotinine-treated vessels (right-shift of all curves vs. CTRL; **B**). Pre-contracted cotinine-treated vessels relaxed normally with acetylcholine (ACh) and sodium nitroprusside (SNP; **C**), but N_PRE_ showed evidence of endothelium-dependent dysfunction. The mRNA expression ratios are shown relative to β-actin. PKC mRNA levels were consistently ∼3-6 fold lower (**D**; overall p<0.025) and adrenergic receptor (ADR) levels marginally higher (**E**; overall p=0.05). Conclusion: adrenergic hypo-contractility develops after birth due to a cotinine-related signalling anomaly affecting two key genes located upstream from the smooth muscle contractile proteins. Means ± SEM.

## Cotinine reciprocally regulates two key genes involved in smooth muscle contraction

PKC mRNA expression was consistently lower (p=0.025; Figure 5D) and ADR mRNA marginally higher (p=0.05; Figure 5E) in vessels from all drug-treated litters (no significant change in expression of other gene targets). This finding suggest that the vascular hypo-contractility caused by nicotine was actually secondary to an intra-cellular signalling anomaly caused by low-level exposure to its metabolite cotinine.

## Discussion

“Dosis sola facit venenum - the dose alone makes the poison” - has been a dictum of toxicology for 500 years (23). Here we extend this principle to cotinine, long neglected and widely regarded as little more than a proxy measure of recent exposure to numerous other potential toxins in tobacco, including nicotine (24). Far from being inert, cotinine in fact has latent, highly selective actions on the immature heart and blood vessels, notwithstanding a normally wide safety margin, by virtue of prolonged tissue retention. Our conclusions are in line with human / rodent studies that show acute exposure *of adults* to this substance relaxes smooth muscle, widens blood vessels, attenuates adrenergic constriction, slows the heart, and dose-dependently lowers BP (8, 9, 25-27). These actions may in fact be so effective in counteracting the sympathetic excitation-hypertension-tachycardia of nicotine as to explain why the BP of habitual smokers is often lower (not higher) than non-smokers (25, 28-30). The surprising, novel findings from our modeling of short-term, *perinatal* exposure to cotinine is that its effects emerge very early-on, at blood levels equivalent to the lowest reported, and represent a legacy of *past* (not recent or ongoing) exposure i.e. cotinine’s cardiovascular consequences endure long after it has disappeared from the environment.

Some natural substances are toxic in small amounts, others harmless except at high doses. Our “stepwise” exclusion strategy – administer both drugs (N_PRE_), withdraw one (C_PRE_), and narrow the period of exposure (C_POST_) – reveals how the onset, duration, and blood level (i.e. cumulative *effective* dose) of a major tobacco by-product can have discrete and/or overlapping developmental effects that dwarf the hypothetical risks of many more potent trace constituents of tobacco. Our key finding: exposure to a low cotinine dose for a short time after birth re-programs cardio-circulatory function; a higher cumulative dose for longer (from conception) has a qualitatively similar impact. The inference is clear: the newborn is exquisitely vulnerable to this substance. Why is uncertain, but xenobiotic actions depend on many things, including maturity and gender (31). Our findings are timely given the global extent of inadvertant maternal-infant exposure to cotinine via multiple overlapping routes (32, 33). The blood of newborns of passive smokers contains ∼3 ng/ml cotinine (34), and urine of infants living with at least one smoker ∼25ng/ml cotinine (1, 35) (urine levels are ∼6-fold higher than blood) (6). This *minimum* average exposure may well understate the net cotinine load in many communities (36). So, our modelling in mice is realistic and may even be conservative i.e. understate cotinine’s true developmental potential. How do parent alkaloid and its metabolite interact? Although not addressed in great detail, we do show that exposure to both from conception can generate a “hybrid” phenotype: classical nicotine-like side effects - on breathing (17, 37) endothelial function, possibly vascular structure - combined with enduring, cotinine-like side effects (vascular hypo-contractility); subtle effects of cotinine - on resting BP and HR - seem to fade or are masked. Interaction effects are unpredictable: our empirical model suggest simply that for nicotine “a little goes a long way”, for cotinine “less is more”.

Cotinine appears to impair vascular tension, reactivity, and tone by down-regulating the activity of an enzyme (PKC) that relays and amplifies messages from the cell surface to the interior, and which is a key player in cell homeostasis and disease pathogenesis (38). Low PKC activity is only one explanation for the loss of adrenergic reactivity we observed (right-shift in vascular reactivity curves and ∼two-fold higher EC50). We do not rule out other deficits in smooth muscle sensitizing pathways, although we found none. PKC has diverse, tissue-specific actions: it promotes actin-myosin binding and vascular smooth muscle contraction, modulates activity of ion channels and pumps, hormone and neurotransmitter release, gene expression, protein and DNA biosynthesis, and cell differentiation, proliferation, migration and growth. Conceivably therefore, altered PKC activity could have other long-term cardiovascular (e.g. remodeling, proliferative, atherogenic) consequences (39, 40). Precisely how cotinine exerts its effect on PKC is unclear. Sustained over-activation by low dietary levels of natural plant toxins can “turn-off” (deplete) signalling-competent PKC (41) and blunt the contractile response to catecholamines (noradrenaline, adrenaline) and adrenergic agonists like nicotine (37, 42). Loss of signal coupling to smooth muscle contractile proteins may initiate a compensatory up-regulation of the membrane (adrenergic) receptors that mediate sympathetic vasoconstrictor / pressor reactivity to “re-balance” the system (43), but this may not fully reverse a hypo-contractile / hypo-adrenergic state or restore BP (Figure 5D-E; Table 1). Of course, mRNA levels are only indicative of transcriptional gene regulation: with limited tissue available we did not show that cotinine down-regulates PKC (and upregulates ADR) proteins. Changes in large elastic conduit vessels (the aorta) are not *necessarily* representative of the microcirculation, however PKC does regulate the calibre of these small resistance vessels, so the mechanism we propose is broadly consistent with cotinine’s ability to lower BP (Table 1) (44). Nicotine’s actions are more extensive and profound by virtue of its strong affinity for a ubiquitous, neuronal nAChR subtype which modulates catecholamine release from perivascular nerves and into the blood stream (17, 37). Its cytotoxicity is also driven by the release of reactive O_2_ species that impair endothelial NO production and weaken arterial dilation-relaxation, and causes smooth muscle cells to proliferate and secrete connective tissue (45, 46). A thicker, stiffer (N_PRE_) vessel may contract more forcefully when treated with an agent that bypasses cell surface adrenergic receptors to directly depolarize / flood the cell with Ca^2+^ (Figure 5A).

The small arteries and arterioles are normally partially constricted at rest; this basal tone reflects a balance between factors that contract and relax the underlying smooth muscle – a process fundamental for matching blood flow to changing systemic and regional demand (47). Functional hyperemia - dilation of local capillary-arteriolar networks and up-stream supply arteries in metabolically active muscles – is the reason why BP falls slightly during systemic hypoxia (48). Cotinine’s effect of weakening basal tone and shifting the prevailing balance towards vaso-relaxation, may explain the lower heart rate and blood pressure observed at rest (Table 1), and also mitigate any local hypoxic vasodilatory effects (Figure 4). A slower heart and longer cardiac filling-time may be partial reflexive compensation for reduced cardiac filling and output (hence a lower resting BP, since the heart only pumps blood it receives). Whether cotinine slows the heart directly or indirectly has not yet been established (9).

The classic connotation of toxicology (“the science of poisons”) is how chemicals and biology interact. With only a fraction of nicotine’s potency, cotinine may be less likely to elicit undesirable side effects, but its actions need careful scrutiny to ensure it is not toxic by other mechanisms. The parallel between the circulatory changes we attribute to cotinine in mice, and the remarkably similar, subtle deficits described in newborns who breathe low-level smoky air for a few months (1) is so striking it suggests an answer to the question we posed at the outset: that circulatory dysfunction in early infancy is plausibly, conceptually linked to low-level, short-term exposure to cotinine. Subtle, underlying deficits in circulatory control during sleep have a variety of important pathophysiologic implications in infancy (49-51). Those for whom these findings are especially relevant are the many millions of pregnant women and their infants regularly, avoidably, exposed to cotinine (mainly via SHS), as well perhaps as advocates of the therapeautic virtues of NRT during pregnancy and/or breastfeeding as a quit-smoking strategy (14, 15). Is it really “safer” to administer pure nicotine to a pregnant mother as a strategy to avoid tobacco’s numerous other impurities? Is it less harmful to consume one chemical which has significant circulatory side effects inextricably linked to its principal metabolite?

## Conclusions

Cotinine is far more than simply an inert biomarker of recent perinatal exposure to tobacco and tobacco smoke: some side effects of nicotine are in fact plausibly mediated by (liver) conversion to this substance. Our findings suggest that the goals of all adult smoking cessation strategies should be broad: to reduce, preferably to eliminate completely, all fetal and infant exposure to nicotine *and* its metabolites. Cells reprogrammed - by cotinine as well as by nicotine - may never completely revert to their original state: “All changed, changed utterly: a terrible beauty is born” (52).

## Clinical Significance

Cotinine is used routinely to gauge recent perinatal exposure to tobacco smoke but has been of no health concern itself until now. Here we show that short-term, low-level exposure to cotinine alone before or after birth reprograms cardiovascular function. These cardiovascular effects endure long after cotinine has completely disappeared from the environment. The actions of cotinine appear to be the reverse of nicotine: it relaxes blood vessels and weakens vascular tone and blood pressure regulation during sleep. A selective, real-life synergism may exist between both substances. Advocates of nicotine replacement therapy as a quit-smoking strategy (a “necessary evil”) during pregnancy and breastfeeding must now consider an additional associated risk: that the developmental toxicity of nicotine is in part inextricably linked to its main break-down product in the body, cotinine.

## Acknowledgments

None

## Author Contributions

GC designed the study; SB, VL-M, CB, AS, GZ, HL, AA, GC performed the research and analyzed data:; GC wrote the paper: all authors revised and approved the manuscript.

